# A mechanical parameter of coasting varies bout to bout and predicts swim timing in larval zebrafish

**DOI:** 10.64898/2026.09.25.754473

**Authors:** Javier J How, Sina Schwinn, Martin Haesemeyer, Florian Engert

## Abstract

Most animals move intermittently. Of these, animals that fly and swim move in two stages: during active propulsion, when they beat their wings or swing their tails, and right after, when they glide or coast through the air or water. The kinematics of active movements are typically well studied, but the coasting period afterwards is less understood. We used larval zebrafish as a model of intermittent locomotion. Larval zebrafish swim in discrete bouts lasting ∼200 ms, coast for short durations and distances, and then are largely stationary for ∼800 ms. We find that larval zebrafish do not coast like a passive object. While they coast for distances that are correlated with the prior swim’s strength, as expected from physics, they seem to be able to modulate the mechanics of their coasting: the time constant is not a constant, but varies from coast-to-coast. Furthermore, the time constant is correlated with the duration of the interval between swim bouts. This suggests that larval zebrafish can either shorten or extend their coasting duration, and that coasting inhibits the initiation of the next swim. Finally, the time constant changes quickly, while other swimming kinematics (e.g., distance swum) are autocorrelated over the course of at least several seconds. Altogether, this suggests that fish, and perhaps other animals that move intermittently, actively modulate their post-propulsion movement, possibly to gather sensory feedback in the absence of blurry visual feedback associated with fast locomotion.

## Introduction

Understanding behavior requires knowing which of its features are controlled by the nervous system, and which follow from the body and its physical environment. This is usually hard to determine because motor commands and body mechanics act simultaneously, so any movement reflects both at once. But it matters for two reasons: a feature assumed to be under control invites a search for a neural signal that may not exist, while a feature assumed to be mechanical may vary in ways no one thinks to look for.

Larval zebrafish are intermittent swimmers, alternating brief bouts of tail beating with longer passive intervals. Work on this behavior has largely treated the cycle as two states – an active bout and a pause – and focused on the former (Dunn et al., 2016; Johnson et al., 2020; Marques et al., 2018). But the fish does not stop moving when the tail does. Momentum from the bout carries the fish forward, and the early part of the “pause” is in fact occupied by a glide. The mechanics and ontogeny of this process, called coasting, have been previously studied: McHenry and Lauder (2005) found that larvae and juveniles coast at a low Reynolds number and therefore in a viscous regime, where drag is linear in velocity and the coast decays exponentially, with a time constant set by body mass and drag. Conversely, they found that adults coast at a high Reynolds number and therefore in an inertial regime, where drag is quadratic in velocity and the coast decays far more slowly, carrying the fish a much greater distance relative to its body length. This work explored how coasting changes across development but did not examine if it affects ongoing behavior.

Here, we study the dynamics of coasting and its effect on subsequent behavior by examining 41,635 swim cycles across 139 fish. We find that, as expected from physics, the distance the fish coast – the coasting distance – is strongly correlated with the velocity of their preceding swim. Unexpectedly, we find that, when the coast decays more slowly, fish initiate the next swim after even longer pauses, consistent with residual self-motion gating swim initiation. Furthermore, we find that the rate at which a coast decays is not fixed within an animal, as the mechanics of a passive coast predict it should be: it varies far more from bout-to-bout than measurement error or drag-regime effects can account for, and no kinematic or postural variable we measured explains it. Finally, we find that several kinematic features of both the active swimming and coasting periods are autocorrelated for 12 or more bouts, suggesting that fish slowly transition between states, in agreement with prior work (Krishnan et al., 2025; Mangalwedhekar et al., 2026). Altogether, this work provides a basis for understanding the link between two aspects of fish behavior – swimming and coasting – and implicates the sensory consequences of self-motion in the timing of the next bout.

## Results

### Coasting distance is controlled by swim velocity, and the next swim is suppressed during coasting

We examined the coasting dynamics of larval zebrafish swimming for 30 minutes in the dark (Figure 1A). We first extracted several behavioral traces for 139 fish: the angle between the tail tip and a line between the swim bladder and the center between the eyes (i.e., tail angle in degrees), heading (in degrees), and position (in pixels). We then detected swim bouts by looking for local increases in swim vigor (i.e., the 50-ms rolling window standard deviation of Tail Angle; swims are shaded grey in Figure 1B). This allowed us to segment the swim cycle into three periods: active swimming (when the tail is moving and the fish is propelling itself), coasting (when the fish briefly glides but is not moving its tail), and a stationary period (when the fish is effectively still). To analyze the dynamics of the coasting period, for each interbout interval (IBI; period between bouts) following each swim we computed the fish’s distance from the start of the IBI, dist_origin_ (Figure 1C, bottom panels). Coasting in the viscous regime can be modeled as an exponential decay in velocity or, equivalently, an exponential rise in dist_origin_ (McHenry & Lauder, 2005). We modeled dist_origin_ as a rising single exponential with 3 parameters (Equation 2b): the total coasting distance *A*, the time constant *τ* (which measures how quickly velocity decays), and the offset *C* which captures some tracking noise. These parameters and fits are shown as the green curves in Figure 1B for three coasts from an example fish (fish #129): a short, medium, and long coast.

**Figure 1.**
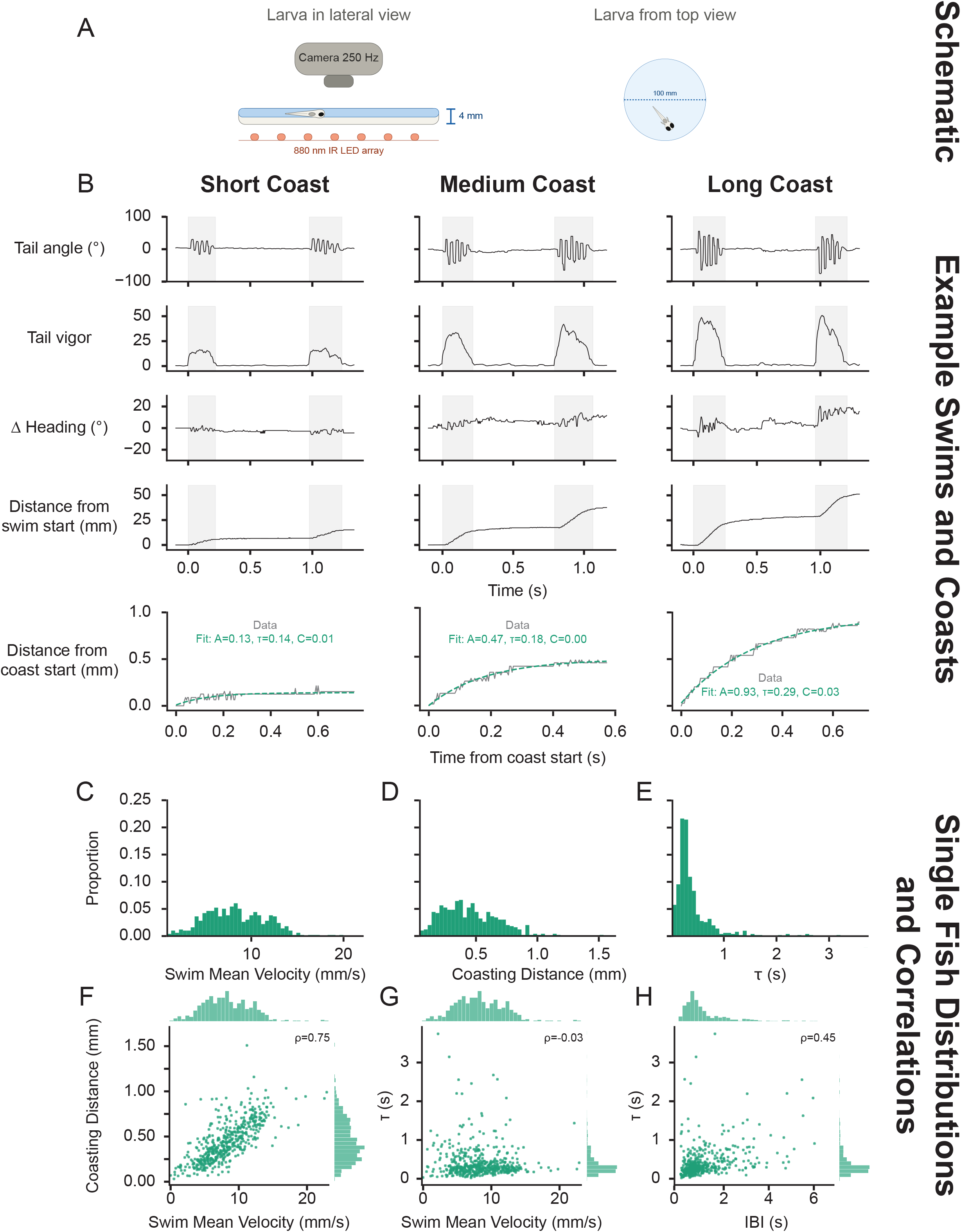
Coasting in a single fish. (A) Experimental setup. (B) Example traces of tail angle, tail vigor, heading, and distance from location at swim start. Shading indicates period when swims were detected. Coasting period is separately plotted for the IBI, with overlaid fit (green), below; distance is from location at coast start. (C – E) Distributions of (C) mean swim velocity, (D) coasting distance, and (E) *τ* for this fish. (F – H) Correlations, for this fish, between (F) mean swim velocity and coasting distance, (G) mean swim velocity and *τ*, and (H) interbout interval duration (IBI) and *τ*. The green histograms along the axes represent the spread. n = 508 traces in C – H.

Larvae inhabit a linear drag regime (McHenry & Lauder, 2005), which makes three predictions that have not previously been tested. We first tested these predictions in 1 example fish (#129). First, the coasting distance *A* should be correlated with the velocity of the fish upon entering the coast, *v*_0_. In agreement with this prediction, we found that the mean velocity during the swim (median 8.1 mm/s, IQR 5.8–11; Figure 1C) – a proxy of *v*_0_ – is strongly correlated with how far the fish coasts (median 0.43 mm, IQR 0.28–0.62; Spearman rho = 0.75; Figure 1D, F). Second, the time constant *τ* (median 0.29 s, IQR 0.20–0.45; Figure 1E) should not be correlated with *v*_0_ or mean velocity; our results for fish 129 agree (Spearman rho = -0.03; Figure 1G). Finally, *τ* should be independent of the duration of the IBI (median 1.02 s, IQR 0.78–1.41 s; Supplementary Figure 1A) because *τ* is a function of the fish’s mass and drag coefficient (Supplementary Note), neither of which depends on IBI. Surprisingly, however, we find that *τ* is correlated with the duration of the IBI (Spearman rho = 0.45; Figure 1H).

These findings were consistent across all 139 fish (Figure 2D – F; green dots are data from fish 129, grey dots from all other fish), and robust to filtering traces (Supplementary Figure 2). Over all bouts, the Spearman rho between swim mean velocity (median 8.4 mm/s, IQR 5.7-11; Figure 2A) and coasting distance (median 0.45 mm, IQR 0.27-0.66; Figure 2B) was 0.75 (Figure 2D); between swim mean velocity and *τ* (median 0.33s, IQR 0.22-0.49; Figure 2C) was -0.03 (Figure 2E); and between *τ* and IBI (median 0.91 s, IQR 0.62–1.34 s; Supplementary Figure 1B) duration 0.46 (Figure 2F). We also computed the Spearman’s rho between each of those parameters per fish and found that the correlations held: the distribution of rho between swim mean velocity and coasting distance *A* had a median of 0.704 (Figure 2G), swim mean velocity and *τ* had a median of -0.044 (Figure 2H), and *τ* and IBI duration had a median of 0.410 (Figure 2I). Both analyses showed that the first two predictions agreed with the physics expected from a passively coasting object, while the third, unexpected finding suggested that coasting seems to prevent the fish from initiating the next swim bout.

**Figure 2.**
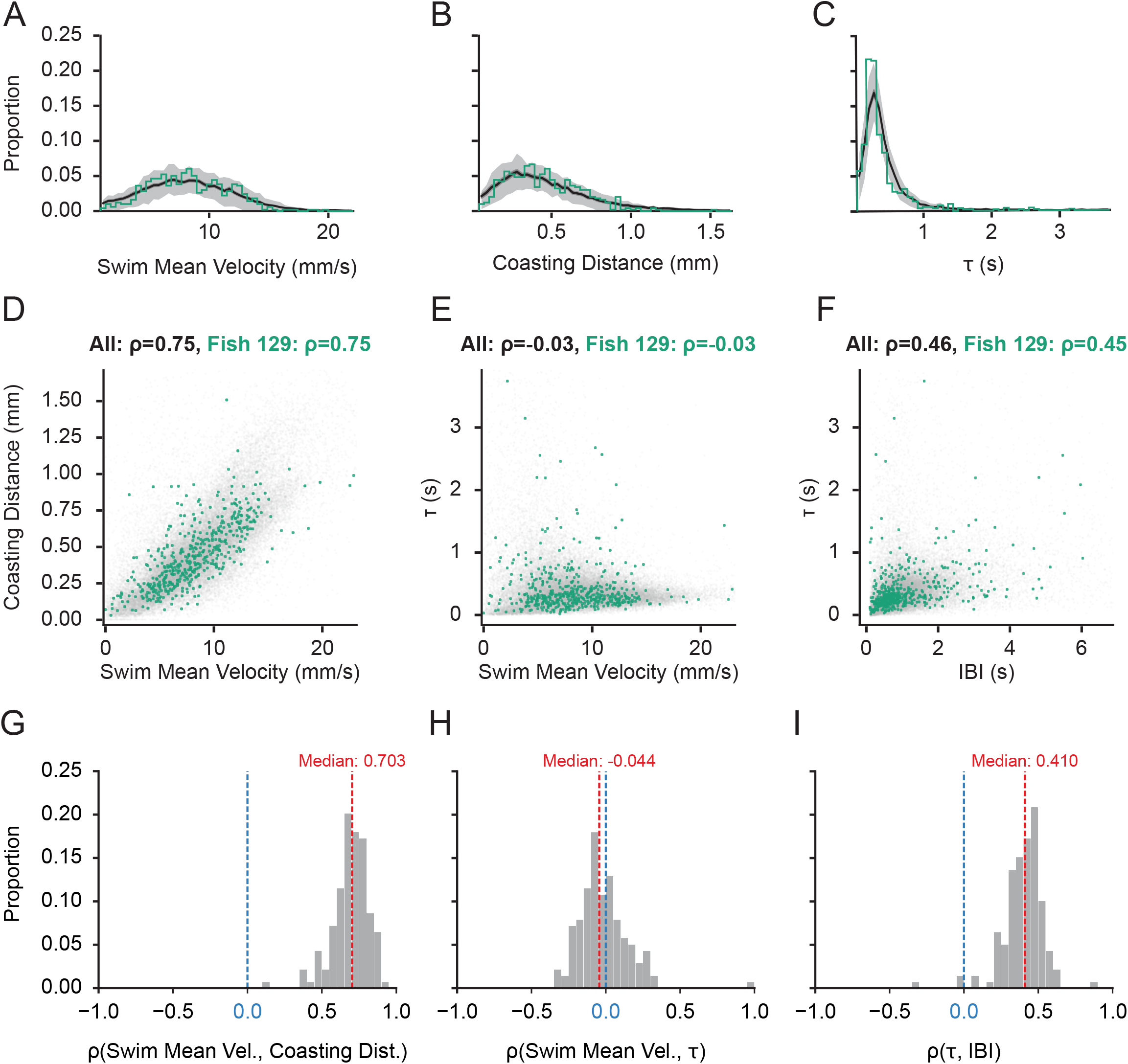
Coasting distance is correlated with swim velocity, and the time constant of coasting is correlated with the duration of the interbout interval. (A – C) Distributions of (A) mean swim velocity, (B) coasting distance, and (C) *τ*, averaged over all fish; black lines are mean, shading is 20th – 80th percentile, and green line is example Fish 129 from Figure 1. (D – F) Correlations, for all fish, between (D) mean swim velocity and coasting distance, (E) mean swim velocity and *τ*, and (F) IBI and *τ*; Fish 129, from Figure 1, is in green. (G) Histogram of correlation coefficients between coasting distance and mean swim velocity for all fish. (H) Histogram of correlation coefficients between mean swim velocity and *τ* for all fish. (I) Histogram of correlation coefficients between IBI and *τ* for all fish. All correlation coefficients are Spearman’s rho. n = 139 fish, 41635 traces.

Altogether, we find that the active swimming period sets the coasting distance, while the time constant of the velocity decay during coasting affects subsequent swim initiation.

### Time constant *τ* varies more from coast-to-coast than expected from a passive coasting process

The time constant *τ* is equal to *m*/*b*, where *m* is the object’s mass and *b* is the object’s viscous drag coefficient. Neither of these properties change over the course of a 30-minute session. However, the distribution of 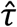 (*τ* values from fitting single exponentials) we get from fitting the coasts of fish 129 is not a delta function, but a wide distribution with a sharp peak at low values of 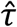 and a long tail (Figure 1E). This was true of all fish, as seen in the averaged histogram in Figure 2C. The wide distribution of 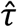 may be caused by two factors: 1) error in our physics model (we model a viscous regime, but an intermediate one is possible), or 2) the fish modulate their 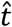 from coast-to-coast.

The first explanation is that the physics of a coasting zebrafish larva is not best described by linear drag (i.e., a viscous regime), nor by quadratic drag (i.e., an inertial regime; likely because they’re too small) (McHenry & Lauder, 2005), but by a combination – that is, an intermediate regime. In this regime, the fish’s velocity when entering the coast dictates the ratio of inertial to viscous drag, with higher velocities leading to inertia dominating early in the coast (see Supplementary Note). This would change the shape of dist_origin_ and so, even with a fixed *τ*, fitting a single-exponential would produce a distribution of 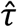 that reflects variability in *v*_0_. To test this, we repeated the simulation with the intermediate model, holding both *τ* and *α* (the ratio of inertial to viscous drag coefficients) constant within each fish. We estimated *α* to be 0.0123 s/mm (Supplementary Note). Each coast’s *v*0 was then obtained by inverting the model from its observed amplitude *A* (see Supplementary Note), so the simulated coasts match the observed distribution of coasting distance.

We found that, for example fish 129, the distribution of 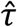 from the simulation (Figure 3A, red distribution) is not as wide as the empirical distribution (Figure 3A, green distribution; the width was computed as the interquartile range of log 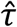). The median width ratio (width of observed distribution divided by width of simulated distribution) was 5.5 (IQR 4.0–8.2), and exceeded 1 in every fish (minimum = 2.1, n = 132; Figure 3B). Aside from having a wider distribution of *τ*, the empirical distribution also has up to fivefold more coasts with a short or long *τ*, with correspondingly fewer at the median (Figure 3C; adjusted p < 0.05 by Wilcoxon test with Bonferroni correction). This indicates that fish can shorten (smaller *τ*) and extend (larger *τ*) their coasts.

**Figure 3.**
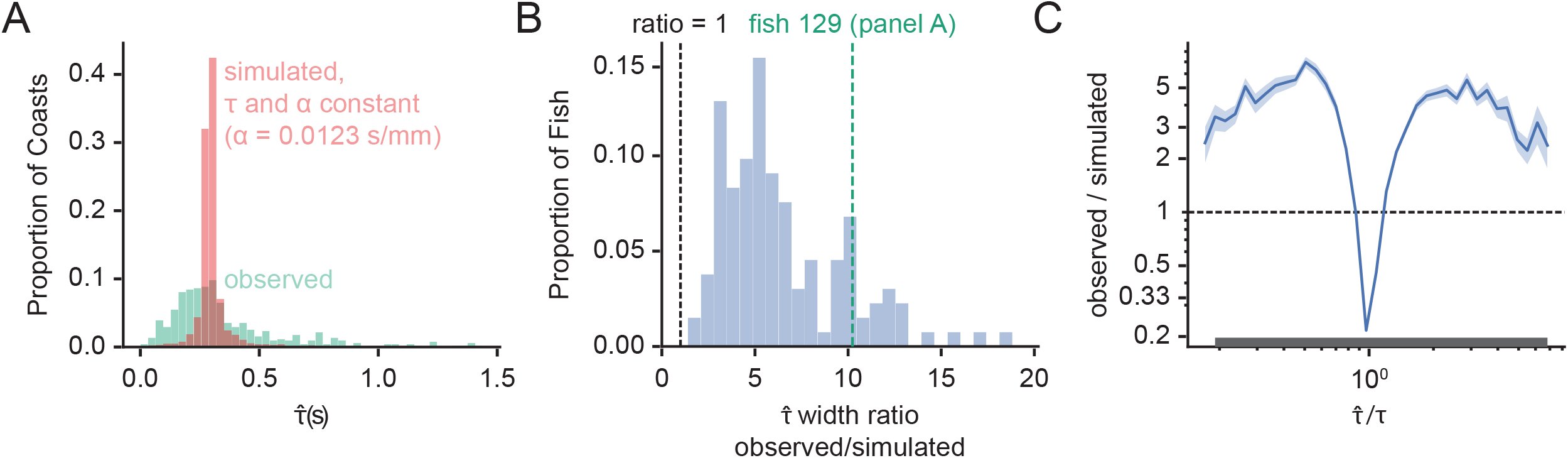
Smaller and larger *τ*’s are more common than expected if the mechanical parameters were fixed. (A) The distribution of the fit *τ* (green) – 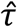 – is wider than what is expected under simulations of the intermediate regime where both *τ* and *α* are held constant (red). For the simulation, *τ* was set to the median for Fish 129, and *v*0 for each simulated coast was obtained by inverting *A* = (*τ*/*α*) · *ln*(1 + *αv*0), using that coast’s observed *A. α* was fixed at 0.0123 s/mm for all fish. (B) The distribution of the ratio of the interquartile range of log 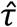 for the real data to that of the simulated 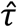. Dashed black line marks a ratio of 1, and dashed green line is the width ratio for Fish 129. (C) The ratio of the observed to simulation histograms for all fish; line is mean, shading is SEM. The grey bars at the bottom are bins where the difference was significantly different from 0 by Wilcoxon test with Bonferroni correction for the number of bins (number of bins = 61).

The remaining explanation is that *τ* changes from coast-to-coast. *τ* is a function of mass and the drag coefficient (Supplementary Note); while the former should not appreciably change in a 30-minute-long session, the latter *could* vary if the fish changes its posture to become more streamlined. We reasoned that the fish could change its drag by 1) moving its tail, which we can measure, or 2) repositioning its pectoral fins, which we cannot. We tested the first directly. When fish begin to swim, they swing their tails to a maximum of 57±1° (mean ± SEM across fish, n = 139; Figure 4A), their tail vigor rises to 36±0.7° (Figure 4B), speed peaks at 9.8±0.2 mm/s (Figure 4C), and the tail beat frequency reaches 21±0.2 Hz with a max amplitude of 46±0.9° (Figure 4D). These are the markers of tail movement that are associated with active locomotion. By comparison, by the time the fish begins to coast, all of these kinematic features have fallen to the noise floor of each measure (Figure 4E – H), such that we are not able to detect any tail beating (Figure 4H, the frequency cannot be estimated where the amplitude goes to 0). A sustained posture also could not account for the variation in *τ*: neither mean tail angle nor mean tail curvature during the coast was correlated with *τ* across fish, nor was heading variability during the coast (Figure 4I). While swim displacement, mean velocity and vigor predicted coasting distance strongly (Figure 4K), these features predicted nothing about *τ* or IBI duration (Figure 4I, J), so the null results are not attributable to insensitivity of the measure.

**Figure 4.**
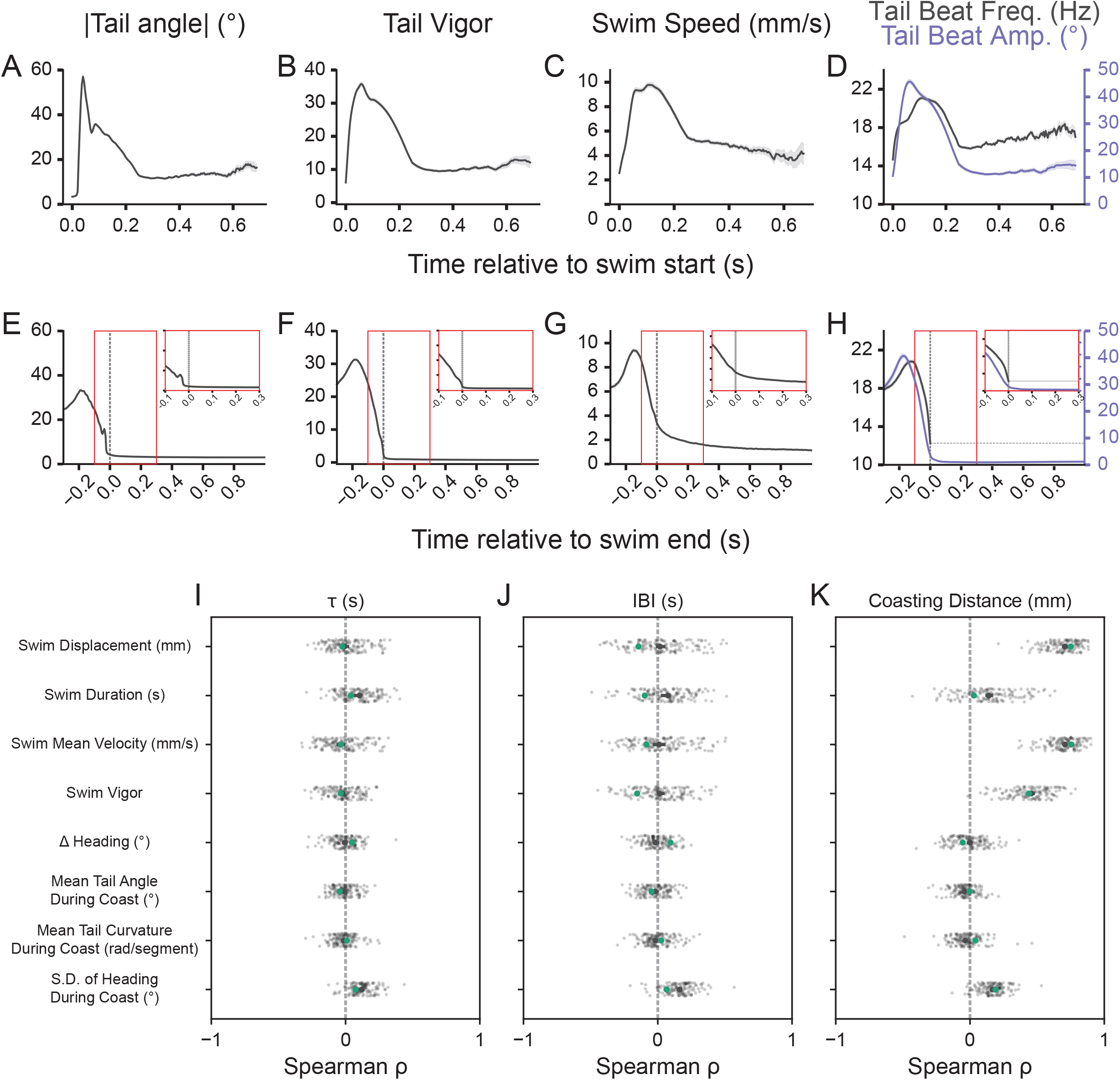
Fish decelerate during the swim and passively coast, and *τ* varies from coast-to-coast independently of measured kinematic features. (A, E) Mean absolute tail angle, (B, F) tail vigor, (C, G) swim speed, and (D, H) instantaneous tail beat frequency (black) and amplitude (purple) locked to (A – D) swim onset and (E – H) swim offset. Vertical dashed grey line in E – H indicates swim offset. The horizontal purple line in H indicates the last frequency before tail beat amplitude dropped below threshold – that is, when the fish stopped moving its tail. Insets in E – H have the same y-axes as the parent panel, but are constrained between -0.1 and +0.3 seconds from swim offset (the red box). (I, J) Variability in (I) *τ* and (J) IBI is not predicted by measured kinematic features. (K) Variability in coasting distance is predicted by swim displacement, mean swim velocity, and swim vigor. Green dots in I – K are Fish 129. Lines in A – H are mean, shading is SEM.

Since the fish does not seem to bend its tail during coasting, we hypothesize that the fish may use its pectoral fins to modulate *τ*. However, the pectoral fins were not tracked in this dataset, so we could not test this possibility, and leave it for future work.

### Time constant *τ*, unlike other kinematic features, is only weakly autocorrelated

Larval zebrafish behavior is autocorrelated over time. For instance, their performance on a sensorimotor task varies over time, partly due to drifting attention and recent stimulus history (Krishnan et al., 2025; Mangalwedhekar et al., 2026). They also string together several (usually 5 or more) turns in the same direction when swimming in a largely homogeneous environment (Dunn et al., 2016). It is unclear, however, how a drifting behavioral state affects coasting behavior, or other kinematic features of swim bouts.

We first found runs of consecutive swim bouts and coasts that were reliably tracked, keeping a bout and its subsequent coast only if both were well tracked (see Methods). Within each fish, we then computed the autocorrelation of several kinematic features across bouts. A given lag was reported only if at least 10 fish contributed at least 10 bout pairs each at that lag, which limited the analysis to lags of 12 bouts or fewer. *τ* was the least autocorrelated of the variables measured (0.075 at lag 1, falling to 0.02 by lag 5; Figure 5A). Coasting distance and IBI duration were more strongly autocorrelated (0.22 and 0.17 at lag 1, respectively; Figure 5B, C), and coasting distance remained elevated at ∼0.15 across all 12 lags. Among swim features, mean velocity and displacement were the most persistent (0.28 and 0.22 at lag 1, respectively, remaining near those values out to lag 12), while vigor and duration were weaker (0.09 and 0.11, respectively) and fell to near zero by lag 12 (Figure 5D–H). Change in heading decayed to near zero within 3–5 bouts, consistent with prior work (Dunn et al., 2016; Figure 5I).

**Figure 5.**
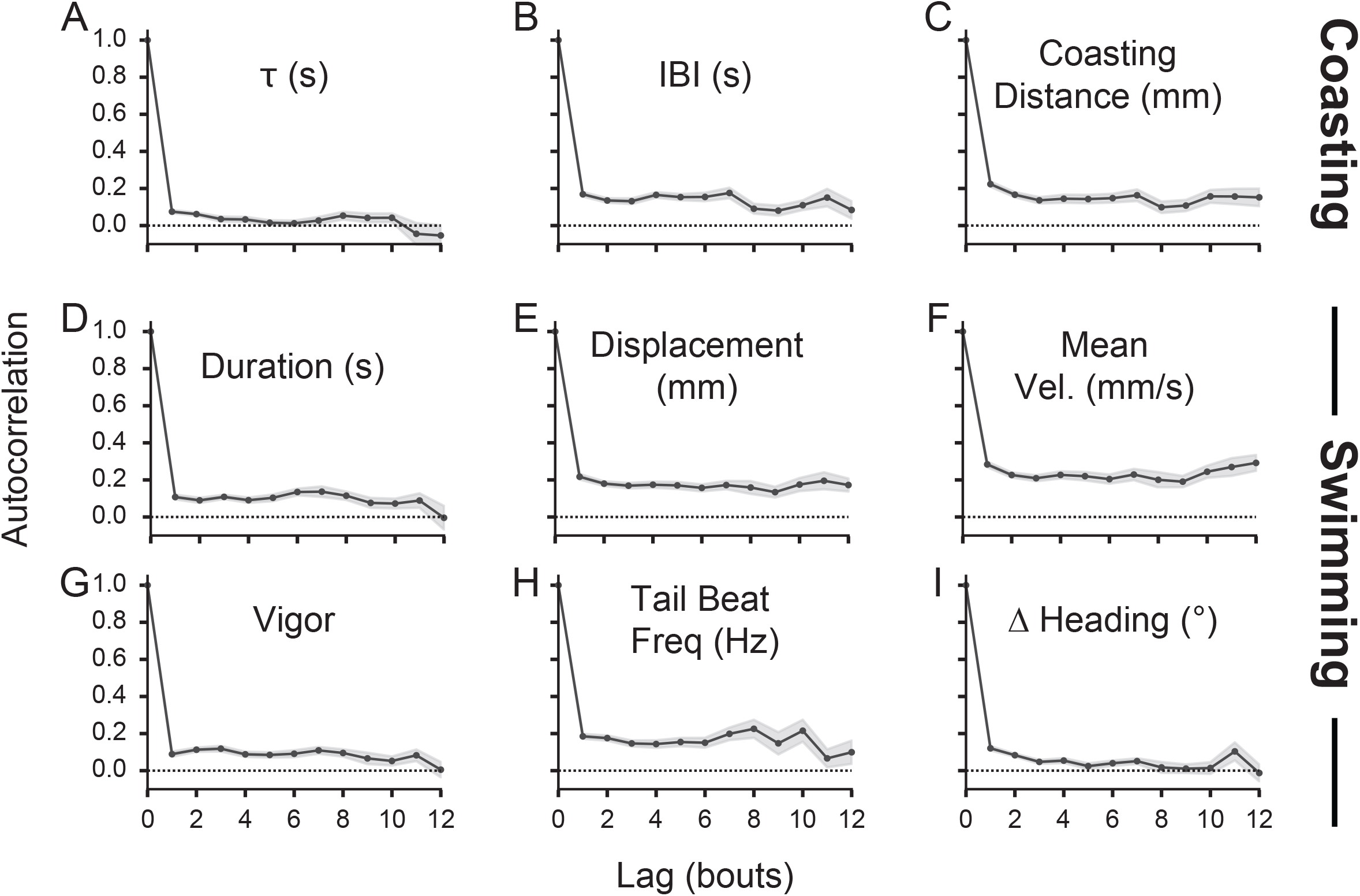
*τ*, unlike most other kinematic features, is weakly autocorrelated. (A – I) Autocorrelation, across bouts and over all fish, of several coasting features: (A) *τ*, (B) IBI, and (C) coasting distance; as well as active swimming features: (D) duration, (E) displacement, (F) mean velocity, (G) vigor, (H) median tail beat frequency, and (I) change in heading during the swim. Lines are mean, shading is SEM.

## Discussion

We propose that the swim cycle of larval zebrafish is best divided into 3 parts: active swimming, coasting, and stationarity. We examined the physics of coasting to show that 1) the distance coasted is correlated with the strength of the preceding “active swimming” period, as expected from the physics of a passively coasting object, but that 2) the time constant of coasting, *τ*, is correlated with the duration of the interbout interval (IBI), which is not expected from a passively coasting object. This suggests that coasting is an active process, not governed solely by the physics governing how velocity decays in water (McHenry & Lauder, 2005), but under neural control.

### How do fish modulate *τ*?

The distribution of *τ* in real fish shows that, compared to our simulation, the velocity of some coasts decays more quickly while others decay more slowly than expected (Figure 3). In a way, some coasts end early, and others last longer. We propose that there are two ways to think about this phenomenon: one focused on what the fish does to modulate *τ*, and another focused on how the duration of the coast modulates what the fish does.

First, we focus on the fish as the initiator. Short coasts correspond to situations where the fish chooses to brake, and long coasts correspond to situations where the fish either accelerates or conserves momentum. We tested the idea that a fish might beat its tail during the coast to gently propel themselves during the coast, and that short and long coasts differ in the amount of this propulsion; however, there seems to be no obvious tail-based propulsion during coasting. It is still possible that an analysis with a higher spatial resolution could resolve finer tail movements, but at a resolution of ∼1/8 mm per pixel, we think this is unlikely. Another possibility is that fish use the pectoral fins to either propel themselves, as cichlids do (Guilbeault et al., 2026), or to change their drag coefficient, as suggested by McHenry and Lauder (2006). Drag in the viscous regime increases with wetted surface area (SA), and extending the pectoral fins might sufficiently increase SA as to decrease *τ*. This mechanism agrees with the finding that larval zebrafish tuck the pectoral fins closer to the body when swimming at higher speeds (Green et al., 2011). We did not, however, have the resolution to measure the amplitude (i.e., extent of abduction) of the pectoral fins, and leave that to future work. In all, by modulating the time constant of coasting, the fish can effectively end it early or late, and initiate the next swim after it is complete.

Second, we consider what variation in coasting means for the fish’s own sensory input. Forward motion produces backward optic flow, which suppresses swimming (Naumann et al., 2016), and differences of ∼1 mm/s in forward-moving gratings are sufficient to modulate IBI and swim latency (Severi et al., 2014). A coast beginning at ∼3 mm/s (Figure 4) therefore generates flow within the range that affects swim timing, and *τ* determines how long it persists: at the 25th percentile (*τ* = 0.22 s) of the observed *τ* distribution a coast remains above 1 mm/s for 240 ms, at the median (*τ* = 0.33 s) for 360 ms, and at the 75th percentile (*τ* = 0.49 s) for 540 ms. The variation in *τ* we observe thus corresponds to a 300 ms difference in the duration of self-generated flow, which is a substantial fraction of the ∼1 s IBI. This is consistent with the *τ*–IBI correlation reflecting that the fish withholds the next bout until residual flow subsides.

### Why do fish modulate *τ*?

As a fish swims, it swings its head from side-to-side, and its heading may also be substantially changing. As a result, visual input during the swim is noisy or blurry, and visual sensitivity might decrease, akin to saccadic suppression in mammals (Wurtz, 2008). We propose that a fish would *want* to modulate the time constant of its coast to make use of a cleaner visual signal than what is available during the swim. Specifically, the fish may use movement during coasting to estimate the distance of objects via motion parallax. Motion parallax gives distance because nearer objects sweep across the retina faster than distant ones for the same amount of translation. Because the retinal signal is less noisy or blurry during coasting, and the trajectory during coasting is typically straighter so retinal motion is mostly translational rather than rotational, the fish can use this period to estimate the distance of objects.

### Swimming and Coasting Kinematics May Be Governed Differently By Internal State

Animal behavior is autocorrelated. For instance, both *Caenorhabditis elegans* and larval zebrafish transition between an exploratory state, where they move over large distances, and an exploitative state, where they move in a small region (Flavell et al., 2013; Gray et al., 2005; Marques et al., 2020). Mice and larval zebrafish go through periods of engagement (attention) and disengagement (inattention) (Ashwood et al., 2022; Krishnan et al., 2025; Pettit et al., 2022). Larval zebrafish also string together several (usually 5 or more) turns in the same direction when swimming in a largely homogeneous environment (Dunn et al., 2016). We find that, in addition to autocorrelation in the active swimming period of the swim cycle, two kinematics of coasting are also autocorrelated – namely, coasting distance and IBI duration – but not *τ*. Since both swim strength and, thus, coasting distance are autocorrelated, this suggests that fish transition through “strong” and “weak” swimming states (Figure 5C – G). Curiously, the duration of the IBI is autocorrelated (Figure 5B) but not correlated with swim strength (Figure 4J), which indicates that fish do not rest longer when they swim harder. Furthermore, *τ* is only very weakly autocorrelated (Figure 5A), even though it is correlated with the duration of the IBI (Figure 2F, I). This indicates that the fish transitions through states which modulate the kinematics of the active swimming, but not the coasting or stationary, period, and implies that a changing internal state can selectively affect behavioral parameters.

## Methods

### Fish strains and Behavioral Setup

Freely swimming experiments were performed according to (Anderson et al., 2025) using 5 – 7 days post fertilization pigmented offspring of incrosses between mitfa^+/-^; Elavl3-H2B:GCaMP6s animals. Sex cannot be determined at these ages.

Fish swam in the dark at a temperature of 21°C for 30 minutes in a 100 mm diameter circular arena with a depth of 4 mm. Video was acquired at 250 Hz, while the arena was illuminated from below with 880 nm IR LEDs. The fish’s position (i.e., centroid of its head), heading angle, and positions of 20 nodes along the length of the tail were extracted using custom code available on Github (Anderson et al., 2025). Paramecia were present in the dish for 11 fish.

## Data Analysis

### Swim Characterization

We computed swim vigor as the 50-ms rolling window standard deviation and detected swim bouts as periods when the vigor exceeded a threshold of 5°, then widened the window to include the closest frames when vigor exceeded 3°. For each swim, we computed 1) swim displacement (distance traversed from the first to last timepoint in the swim), 2) swim duration, 3) swim mean velocity (displacement / duration), 4) swim vigor (integral of the vigor curve), 5) Δ heading (change in heading from the first to last timepoint in the swim), and 6) tail beat frequency (median of the instantaneous tail beat frequency during the swim). The instantaneous tail beat frequency was computed by filtering the tail angle trace at 10 – 30 Hz using a 3^rd^-order, zero-phase butterworth filter, applying the Hilbert transform, and differentiating the unwrapped phase of the resulting analytic signal; these used scipy.signal.butter, sosfiltfilt, and hilbert. The instantaneous tail beat amplitude was taken as the magnitude of the same analytic signal. The instantaneous swimming speed was computed from the tracked centroid using a Savitzky–Golay filter (44-ms window, 2nd order) to take the first derivative of x and y, and the speed as the magnitude of the resulting velocity vector.

### Coasting Characterization

To characterize coasting, we focused on the periods between the end of one swim and the start of the next swim – i.e., the interbout interval *k* (IBI_k_). We began with 103,067 coasts, but after filtering out noisy traces (see Single-Exponential Coasting Simulations) we had 44,935 coasts. Coasting can be modeled as a decaying exponential fit on the fish’s velocity trace during the IBI, where *v*0 is the velocity at the start of the IBI, *τ* is the time constant, and C is an offset (Equation 1; see Supplementary Note).

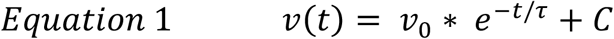

However, because differentiating a noisy position trace to get velocity magnifies the noise, we instead integrated Equation 1 to model coasting as a rising exponential fit on the fish’s distance from origin during the IBI, where origin was defined as its location on the first frame of the IBI (Equation 2a; see Supplementary Note). Here, *A* = *v*_0_*τ* is the total coasting distance (Equation 2b; see Supplementary Note).

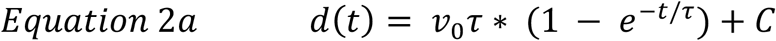

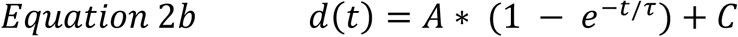

We also fit a ‘flat’ model, with no coasting, where the fish maintains a constant distance (Equation 3). This was used for a filter (see Single-Exponential Coasting Simulations).

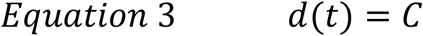

We fit Equations 2b and 3 with SciPy’s curve_fit, with A, *τ*, and C as free parameters. Thus, each coast could be described by its 1) coasting distance A, 2) time constant *τ*, and 3) IBI duration. We then excluded coasts where the estimated *τ* was larger than 10 seconds, as this represented a long tail that was physically unlikely; this excluded ∼7.34% of traces, for a final total of 41,635 coasting traces.

### Single-Exponential Coasting Simulations

We simulated coasting traces to determine if we could successfully recover parameters from noisy traces, and whether the unexpected correlation between *τ* and IBI duration could be due to artifacts introduced by the model fitting or data filtering procedures. We generated 20,000 traces for each of 13 noise levels using Equation 2b, independently sampling *A, τ*, and IBI duration from a range typical of the real fish (i.e., A between 0.01 and 2 mm, *τ* between 0.05 and 1 second, and duration between 0.01 and 3 seconds). We first generated synthetic traces and then added the absolute value of Gaussian noise with σ in [0, 0.01, 0.02, 0.03, 0.04, 0.05, 0.06, 0.07, 0.08, 0.09, 0.1, 0.25, or 0.5].

We then generated various filters to remove extremely noisy or poorly fit traces from our empirical data and tested how their application affected the correlations between our model parameters; ideal filters should remove noisy traces and not induce artifactual correlations in our synthetic data. The filters were: 1) signal-to-noise ratio greater than 3 (SNR, or signal/noise; signal was computed as the 95^th^ – 5^th^ percentile, and noise as the median absolute deviation of the frame-to-frame noise), 2) proportion of spikes less than 1% of the frames of the trace (a ‘spike’ is a frame-to-frame jump exceeding 0.3 mm, and is a tracking glitch), 3) low noise early in the trace (i.e., the second timepoint must be less than 0.1 mm away from origin, and 4) Bayesian information criterion (Equation 4) called it a coasting trace (Equation 2b), instead of a flat trace (Equation 3), or the coasting distance A was less than 0.2 mm (i.e., it was truly a flat trace).

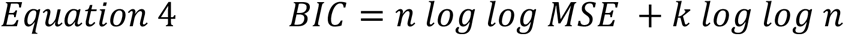

Here, *n* is the number of data points in the trace, *MSE* is the mean squared error of the residuals, and *k* is the number of parameters (0 for flat, 3 for exponential rise).

We found that we could recover the parameters well. At noise sigma of 0 mm, the Spearman rho between our fit *A* (Supplementary Figure 3A) and *τ* (Supplementary Figure 3E) and their true values was 1 for both. The filters did not affect our parameter recovery (Supplementary Figure 3B, F). At noise sigma of 0.1 mm, the Spearman rho between our fit *A* (Supplementary Figure 3C) and *τ* (Supplementary Figure 3G) and their true values were 0.864 and 0.800, respectively, and filtering increased the correlations to 0.920 (Supplementary Figure 3D) and 0.870 (Supplementary Figure 3H), respectively. Higher noise levels did not match the Gaussian noise in the dataset. Thus, we could reliably recover the parameters underlying our single exponential model of coasting.

We found that, across all noise levels tested, the correlation between *τ* and IBI duration on the synthetic traces reached a max of 0.26 Spearman’s rho (Supplementary Figure 4B; examples in Supplementary Figure 3M – P), which was much lower than the 0.46 correlation of the empirical dataset (Figure 2F, I). To test if this result held up under a more realistic noise model, we also generated, for each fish, its own pool of 50 synthetic datasets, where each dataset directly sampled the frame-to-frame noise from the empirical data, and thus captured tracking glitches and other noise sources not found in our Gaussian synthetics. We then compared each fish’s true correlation against that of the 50 null datasets. In 107 out of 136 fish (the subset of 139 fish with at least 20 coasts), the empirical Spearman’s rho was larger than the 95^th^ percentile of the rho from the null datasets (Supplementary Figure 4D), suggesting that the correlation between *τ* and IBI duration is not an artifact, but instead captures a real aspect of fish behavior. By comparison, the correlation in the empirical dataset between A and *τ*, expected from Equation 2b, was similar to that in the synthetics with Gaussian (Supplementary Figure 4A; examples in Supplementary Figure 3I – L) and real noise (Supplementary Figure 4C). Finally, the correlations in the empirical dataset (Figure 2, with filtering) were qualitatively the same without filtering (Supplementary Figure 2).

### Intermediate Regime Coasting Simulations

Larval coasts may span the transition between viscous and inertial drag, which can be modeled together as

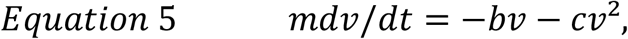

with *τ* = *m*/*b* and *α* = *c*/*b*. This has the closed-form solution

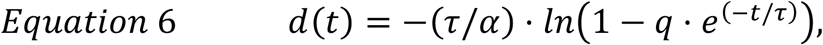

with

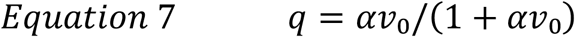

and

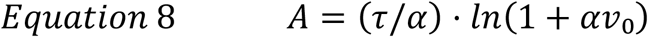

(See Supplementary Note for derivation).

We held *τ* and *α* constant per fish and let only *v*_0_ vary. We obtained each coast’s *v*_0_ by inverting Equation 8 from its observed *A* (i.e., coasting distance), so the simulated coasts reproduce the observed distribution of coasting distances. We estimated *α* to be 12.3 s/m (see Supplementary Note).

### Autocorrelation Analysis

We computed the autocorrelation on truly consecutive swim bouts, defined as bouts where the end of one IBI fell within 20 ms of the start of the next swim (Figure 4). For lag *k*, this meant that the entire run from bout *i* to bout *i+k* had to be uninterrupted; the largest such *k* we could reach, with at least 10 fish contributing data, was 12. A fish contributed to a given lag only if it had at least 10 qualifying pairs at that lag. Within each fish, we first log-transformed features whose distribution was approximately log-normal (i.e., coasting distance A, *τ*, IBI duration, swim displacement, swim mean velocity, and swim vigor). We then z-scored features using the mean and standard deviation (SD) of all of that fish’s values. The autocorrelation at each lag was the mean product of standardized pairs of swim bouts. Finally, we averaged the per-fish autocorrelation functions across fish.

### Statistics

We measured correlations between features by computing Spearman’s rho (scipy.stats). To assess significance, we performed a Monte Carlo test per fish with at least 20 coasts (n = 136 fish) against a simulated null. For each fish we generated 50 synthetic datasets, each matched to that fish in the number of traces and using that fish’s own residual traces as the noise source, so that tracking noise was preserved. Amplitude, *τ*, and trace duration were drawn independently over the same ranges used for the Gaussian synthetics, so any correlation recovered from these datasets reflects estimation artifact rather than a true relationship. Each synthetic dataset was fitted and filtered identically to the real data, and the p-value was taken as the fraction of synthetic datasets whose rho met or exceeded the empirical value (one-tailed; minimum resolvable p-value = 0.02).

### AI Disclosure

We used Claude (Anthropic PBC; Opus 4.6 and 5, and Fable 5.1) to generate code, derive the equations for velocity and position under different drag regimes, and draft parts of the Methods section. We manually verified all outputs.

## Supplementary Note: Coasting under viscous, inertial, and intermediate drag

A larva that stops beating its tail decelerates under fluid drag alone. Writing the equation of motion as

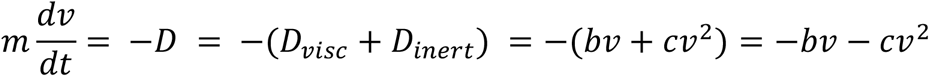

where *D* is the drag force, *D*_*visc*_ is the viscous drag force, *D*_*inert*_ is the inertial drag force, *b* and *c* are the linear and quadratic drag terms, the three regimes correspond to *c* = 0 (viscous; Section 1 below), *b* = 0 (inertial; Section 2 below), and both nonzero (intermediate; Section 3 below). Throughout, *τ* = *m*/*b* and *α* = *c*/*b*, and *x*(*t*) is distance travelled from coast onset, *A* the total coasting distance, and *d*(*t*) = *A* − *x*(*t*) the distance remaining.

### 1. Viscous limit *(c = 0)*

With only linear drag, 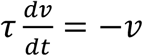

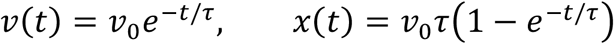

Taking *t* → ∞ gives *A* = *v*_0_*τ*, and hence *d*(*t*) = *Ae*^−*t*/*τ*^. This is the model fitted throughout the paper: a single exponential whose time constant is a property of the body, *τ* = *m*/*b*, and whose amplitude is the product of that time constant with the velocity at coast onset.

### 2. Inertial limit *(b = 0)*

With only quadratic drag, 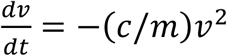, which integrates to

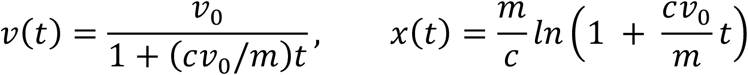

Velocity decays as 1/*t* and position grows logarithmically without bound, so this regime has no finite coasting distance and cannot describe a coast that comes to rest. It is included only as a limiting case; the observed coasts all settle.

### 3. Intermediate regime *(c > 0 and b > 0)*

Dividing by *b*,

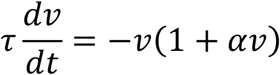

which is separable. Note that this can be rearranged as

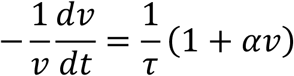

so the instantaneous decay rate of *v* is (1/*τ*)(1 + *αv*); this means that *v* decays at (1 + *αv*_0_) times the viscous rate at onset, settling to 1/*τ* as the fish slows. Thus, inertia changes the rate but does not introduce a second timescale.

Partial fractions give

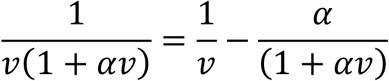

so that integration yields

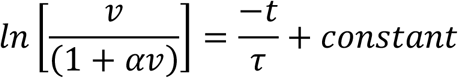

Solving for *v*,

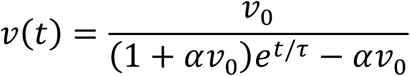

Integrating once more,

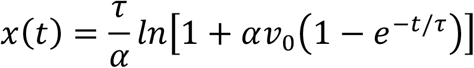

so that, as)*t*→∞,

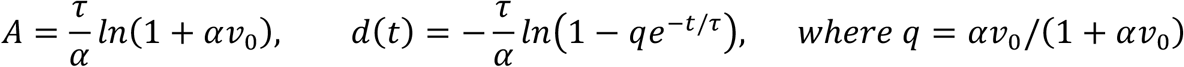

#### Viscous limit

As *α* → 0, *ln*(1 + *αv*_0_) → *αv*_0_ and these reduce to *A* = *v*0*τ* and *d*(*t*) = *Ae*^−*t*/*τ*^, recovering §1 (i.e., the viscous regime).

#### Inversion

Rearranging the expression for *A* (coasting distance),

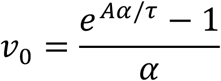

which recovers *A*/*τ* as *α* → 0; in this viscous regime, *A*/*τ* = *v*_0_. This is used to assign each simulated coast the onset velocity *v*_0_ implied by its observed amplitude.

#### Departure from a single exponential

The shape of *d*(*t*) is a function of *q*, which is controlled by the product *αv*_0_, the ratio of inertial to viscous drag at coast onset. Thus, two coasts with the same *αv*_0_ have the same shape, even if *α* or *v*_0_ independently differ; that is, a slow coast for a fish in an inertial regime (low *v*_0_, high *α*) looks like a fast coast for a fish in a viscous regime (high *v*_0_, low *α*). A given fish has a constant *α*, so the shape of their coast varies across bouts due to varying *v*_0_.

The shape of the coasts is captured by

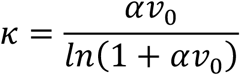

the ratio of the initial decay rate (at coast onset) to the asymptotic rate (as the fish slows); *κ* = 1 indicates that both are equivalent, and the shape is a pure exponential. Conversely, *κ* = 3 means that the coast begins decaying three times faster than it finishes; other *κ* for a range of *αv*_0_ are given in the table below.

| $\alpha v_0$ | 0.2 | 1 | 5 | 10 | 50 |
| --- | --- | --- | --- | --- | --- |
| $\kappa$ | 1.10 | 1.44 | 2.79 | 4.17 | 12.72 |

A single exponential fit to these curves will return a time constant between the two rates, and its position between them depends on *κ*. Since *v*_0_ varies from coast to coast, the fitted 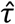 varies even when *τ* and *α* are both fixed, which is the effect exploited in Figure 4.

#### Value of α for larvae

*α* is defined as *c*/*b*, where *b* and *c* are the linear and quadratic drag terms in *D* = *bv* + *cv*^2^, respectively. These are dimensional, and relate to the dimensionless drag coefficients *C*_*visc*_ and *C*_*inert*_ reported by McHenry and Lauder (2005) through their Equations 1 and 2:

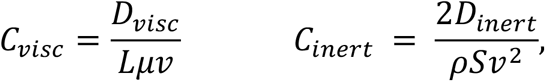

where *D*_*visc*_ and *D*_*inert*_ are the viscous and inertial drag forces, respectively; *ρ* is the density of water, *µ* is the dynamic viscosity of water, *S* is the wetted area of the body, *L* is the body length, and *v* is the swimming speed. We can solve for each drag force in terms of *c* and *b*,

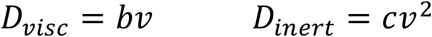

which gives

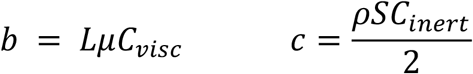

Solving for *α* gives

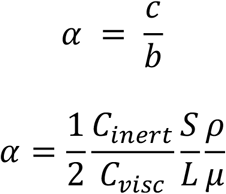

Thus, *α* is a function of the drag coefficients, the morphology of the fish, and properties of the water. We can plug in morphometric values for the average larval zebrafish from Table 1 in McHenry and Lauder (2005), where *S* = 7.89 *mm*^2^ and *L* = 4.35 *mm*. The density *ρ* and dynamic viscosity of water *µ* at 21°C are 998 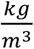 and 9.77 × 10^−6^*Pa* · *s*, respectively.

The values for *C*_*visc*_ and *C*_*inert*_ are 5.72 and 0.076, per Figure 5 and the text of McHenry and Lauder (2005). Each coefficient is a normalization of the total drag, so at intermediate

*Re* (Reynolds number) each is contaminated by the other term – *C*_*visc*_ inflates as the quadratic term grows to dominate, and *C*_*inert*_ inflates at low velocity where the linear term dominates (see Figure 5 of McHenry and Lauder (2005)). Thus, we used values where the fish was purely in the viscous or inertial regimes for *C*_*visc*_ and *C*_*inert*_, respectively. Also note that *C*_*inert*_ is taken from *Re* > 1000, a regime the larvae never reach while coasting; this is not a claim about where larvae operate, but a measurement of a body constant taken where it can be read without contamination from the viscous term.

Thus, *α* ≈ 12.3 *s*/*m*, or 0.0123 *s*/*mm*.

---

**Supplementary Figure 1:**
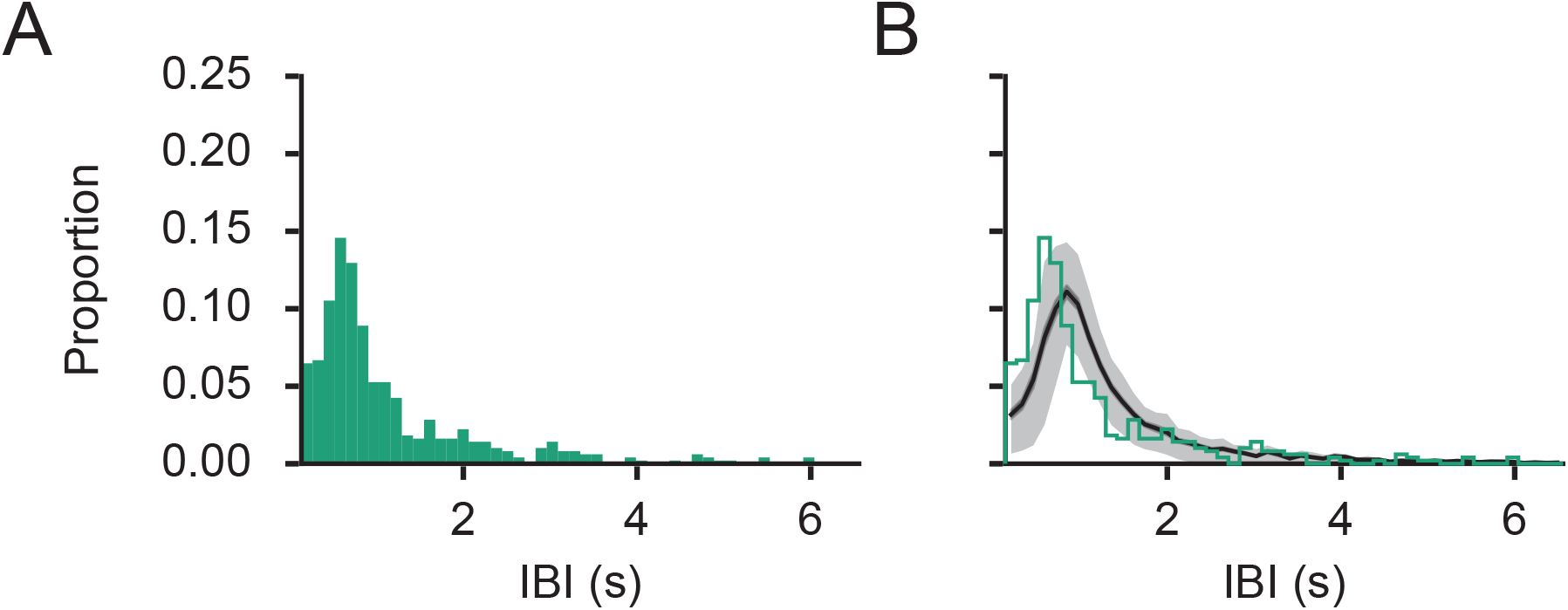
Interbout interval distributions. (A) Distribution of IBI for Fish 129 from Figure 1. (B) Distribution of IBI, averaged over all fish. Black line is mean, shading is 20th – 80th percentile, and green line is Fish 129.

**Supplementary Figure 2:**
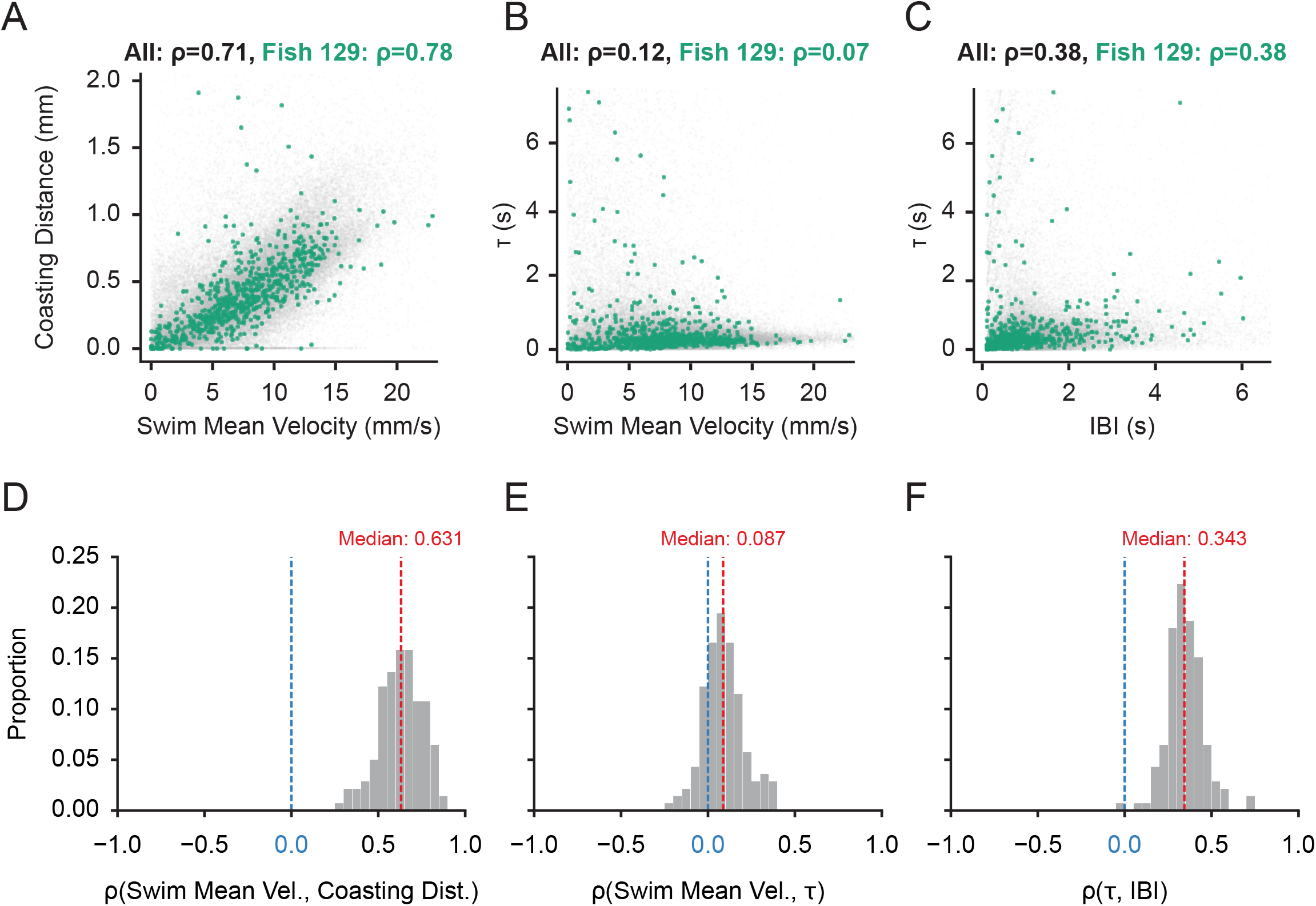
Correlations between features without filtering out traces. (A – C) Correlations, for all fish, between (A) mean swim velocity and coasting distance, (B) mean swim velocity and *τ*, and (C) interbout interval duration (IBI) and *τ*; Fish 129, from Figure 1, is in green. (D) Histogram of correlation coefficients between coasting distance and mean swim velocity for all fish. (E) Histogram of correlation coefficients between mean swim velocity and *τ* for all fish. (F) Histogram of correlation coefficients between IBI and *τ* for all fish. All correlation coefficients are Spearman’s rho. n = 139 fish, 82657 traces.

**Supplementary Figure 3:**
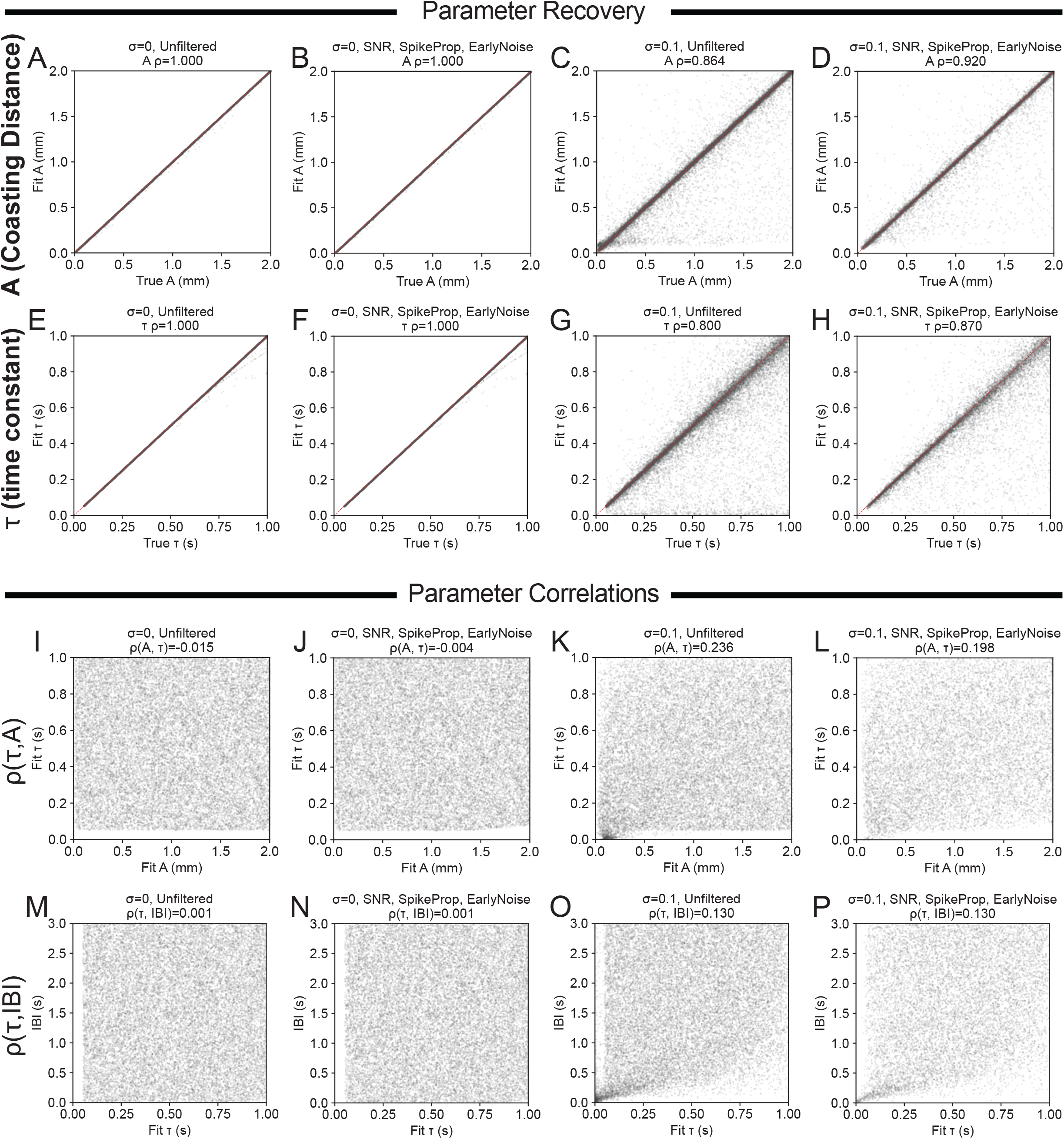
Simulations of coasting with Gaussian noise. Gaussian noise had a standard deviation (σ) of 0 (A, B, E, F, I, J, M, N) or 0.1 (C, D, G, H, K, L, O, P). Data were passed through no filter (A, C, E, G, I, K, M, O) or the full filter (B, D, F, H, J, L, N, P). (A – D) Recovery scatterplots for coasting distance *A*. (E – H) Recovery scatterplots for time constant *τ*. (I – L) Scatterplots of *A* vs *τ*. (M – P) Scatterplots of *τ* vs interbout interval duration.

**Supplementary Figure 4:**
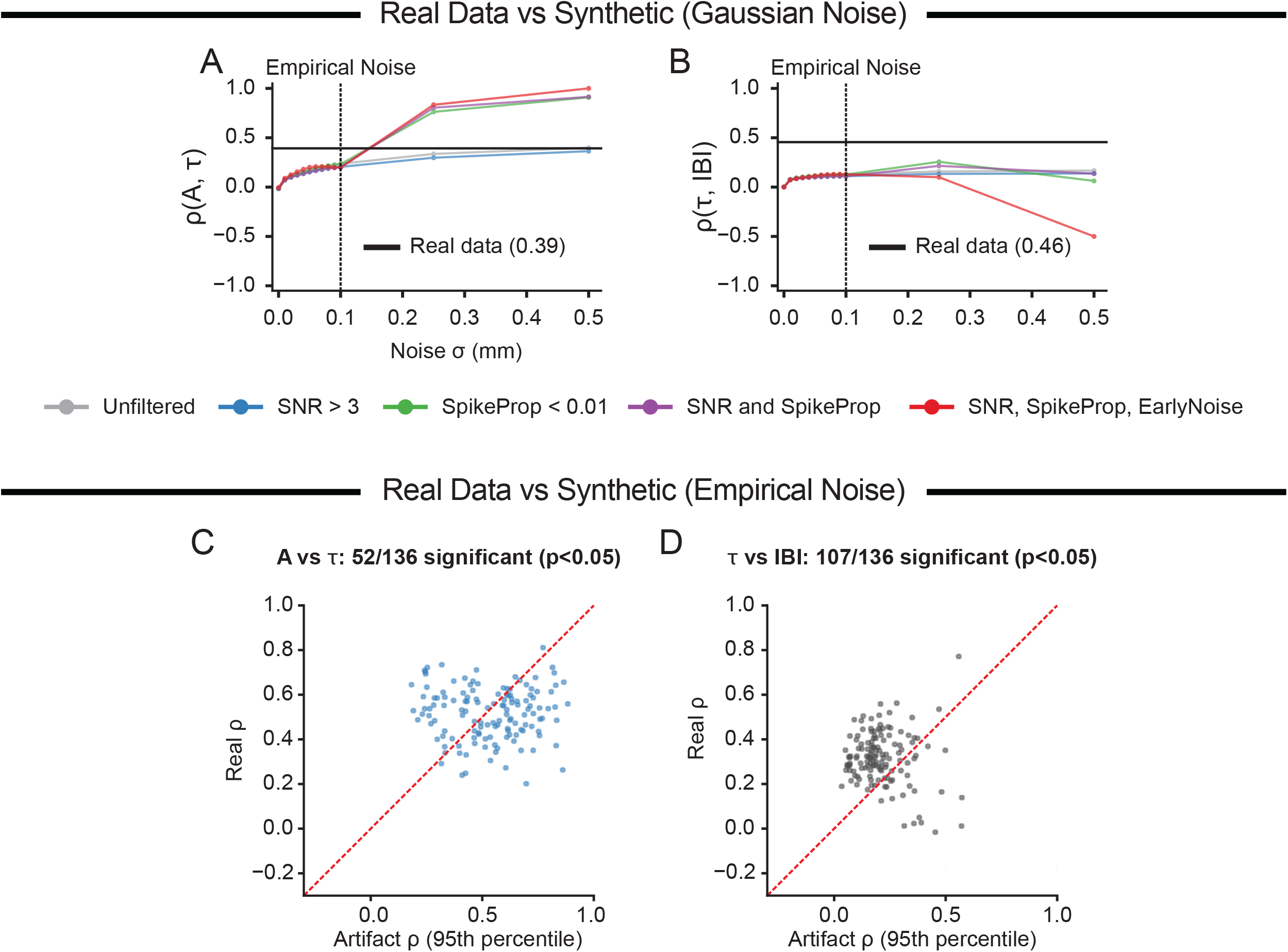
Comparison of correlations between features using real and synthetic data. (A) The correlation between coasting distance (*A*) and time constant (*τ*) in Gaussian synthetics as a function of noise σ and filter applied; real correlation (ρ = 0.39) is thick black line. (B) The correlation between *τ* and the interbout interval duration (IBI) in Gaussian synthetics as a function of noise σ and filter applied; real correlation (ρ = 0.46) is thick black line. (C) Real vs synthetic correlation coefficients between *A* and *τ*. (D) Real vs synthetic correlation coefficients between *τ* and IBI. n = 20,000 synthetic traces for panels A and B. n = 50 synthetic datasets per fish in panels C and D, each with the same number of coasts as the real data. The synthetic correlation coefficients reported in C and D are the 95^th^ percentile coefficient from each fish’s synthetic dataset (i.e., the third largest of 50).

